# Cytokine storm mitigation for exogenous immune agonists

**DOI:** 10.1101/2023.07.07.548130

**Authors:** Irina Kareva, Jana L. Gevertz

## Abstract

Cytokine storm is a life-threatening inflammatory response characterized by hyperactivation of the immune system. It can be caused by various therapies, auto-immune conditions, or pathogens, such as respiratory syndrome coronavirus 2 (SARS-CoV-2) which causes coronavirus disease COVID-19. Here we propose a conceptual mathematical model describing the phenomenology of cytokine-immune interactions when a tumor is treated by an exogenous immune cell agonist which has the potential to cause a cytokine storm, such as CAR T cell therapy. Numerical simulations reveal that as a function of just two model parameters, the same drug dose and regimen could result in one of four outcomes: treatment success without a storm, treatment success with a storm, treatment failure without a storm, and treatment failure with a storm. We then explore a scenario in which tumor control is accompanied by a storm and ask if it is possible to modulate the duration and frequency of drug administration (without changing the cumulative dose) in order to preserve efficacy while preventing the storm. Simulations reveal existence of a “sweet spot” in protocol space (number versus spacing of doses) for which tumor control is achieved without inducing a cytokine storm. This theoretical model, which contains a number of parameters that can be estimated experimentally, contributes to our understanding of what triggers a cytokine storm, and how the likelihood of its occurrence can be mitigated.

## Introduction

Cytokine storm, a life-threatening inflammatory response involving elevated levels of cytokines and hyperactivation of the immune system, has recently gained particular note as one of the causes of morbidity and mortality from coronavirus disease COVID-19 (1). It has previously been observed in a variety of other circumstances, including graft versus host disease (2), bacterial pathogens, (3) and other viral infections, such as SARS (4). Cytokine storms have also been implicated as one of the key culprits in the severity of the 1918 Spanish flu pandemic (5). Additionally, cytokine storms have been observed as a side effect of certain anti-cancer therapeutic interventions, such as chimeric antigen receptor (CAR) T cell therapy (6) and bispecific T cell engagers, also known as BiTEs (7). One of the most notable therapy-induced instances of cytokine storm was the case of a Phase I clinical trial of monoclonal antibody TGN1412, which resulted in severe damage to the health of six volunteers that participated in the trial despite very carefully chosen initial doses (8); numerous additional reports of the details of the case can be found in the literature.

Cytokine storms are most often characterized by severe lung damage, which can lead to respiratory distress, multi-organ failure, sepsis and in some cases, death (6,9,10). Mechanistically, cytokine storms are mitigated by cytokines, which are molecules involved in supporting and regulating the immune response. Cytokines interact in complex networks, geared towards mounting a fast and efficient immune response against pathogens while also preventing excessive damage to normal tissues. If these interactions become destabilized, cytokine storms, or hypercytokinemia, where the immune response causes greater collateral harm than benefit, may occur. Some prominent cytokines that are elevated during cytokine storms include interferon (IFN)-gamma, tumor necrosis factor (TNF)-alpha, as well as interleukins (IL)-6,8 and 10 (1,4,6,9,10). More generally, cytokine storms appear to reflect a scenario in which the immune response itself, rather than a pathogen or immune stimulatory agent, results in pathology. It is this mechanism that this manuscript will explore in greater detail.

Notably, while they are often used interchangeably, there exists a distinction between the terms “cytokine storm” and “cytokine release syndrome” (CRS). Cytokine storm typically refers to an acute reaction, while CRS typically refers to a more delayed response. There exists a discussion about qualitative differences between the two responses, how they are triggered and how they proceed (6), although it appears that the final qualitative dynamics are very similar between the two. Henceforth we will be using the term cytokine storm; however, we believe that the proposed model can be used for better understanding of CRS as well.

One of the most promising types of exogenous immune stimulatory anti-cancer therapies, CAR T cell therapy, can result in a cytokine storm. CAR T therapy requires collecting blood from the patient to isolate T cells, adding a gene into the T cells for a chimeric antigen receptor (CAR) to program these cells to recognize a specific cancer cell antigen, and then introducing the cells back into the patient. This T cell treatment augments the patient’s immunity against their specific cancer. While effective as an anti-cancer treatment with complete remission rates as high as 68 to 93% (11–15), especially in relapsed and refractory hematologic cancer, the risk of a cytokine storm limits the utility of CAR T therapy. Using quantitative methods for predicting and potentially mitigating emergence of a cytokine storm could expand access of CAR T cell and other exogenous immunostimulatory therapies to a larger number of patients.

Several mathematical models have been developed to improve upon our mechanistic understanding of cytokine storm dynamics, and how such storms can be controlled. Waito et al. (16) proposed a thirteen equation mathematical model of cytokine storm, where they grouped cytokines into seven categories based on their pro- and anti-inflammatory properties. They use the model, parameterized with mouse data, to describe the mutual influence of cytokine groups on each other during a cytokine storm. Yiu et al. (17) developed a large scale eighteen equation mathematical model to analyze the data from the TGN1412 clinical trial, using principal component analysis to reveal functional cytokine clusters that were specific to this case. Hopkins et al. (18) created a model of nine major cytokines affecting the outcome of CAR T cell based therapy, and identified two cytokines that, if better-controlled, would theoretically speed up the recovery of patients with CRS. Rana et al. proposed an eight equation within-host model to describe the impact of a cytokine storm on healthy cells in SARS-CoV-2 patients and to study the impact immunomodulatory therapy has on suppressing a cytokine storm (21). These models are all quite complex and high dimensional, aiming to understand how many cytokines, or many components of the immune system, interact during a cytokine storm. A more conceptual model, consisting of only two equations, has been developed by Baker et al. (19) to capture the interactions between pro and anti-inflammatory cytokines in rheumatoid arthritis. It has also been extended into more complex stochastic models in (20) to study conditions under which cytokine storms occur. The complexity of each of these models, with the exception of (19) which does not aim to study cytokine storms, confounds analyzing the broad range of plausible model outcomes in response to infection or treatment with an immune agonist.

Here we propose a conceptual low-dimensional mathematical model that is aimed to capture the general phenomenology of cytokine-immune interactions that may result in a storm-like behavior. We study this in the context of a theoretical model of tumor growth and treatment with an exogenous immune agonist, such as CAR T cell therapy, where drug administration directly affects expansion of cytotoxic immune cells. The model is complex enough to study cytokine-immune interactions, yet simple enough to elucidate the relationship between tumor killing and cytokine storm during treatment. Through model simulations, we ask whether a cytokine storm is an inevitable outcome of a treatment strong enough to eliminate the tumor, or if certain conditions on the dose (amount of drug given) or dosage (duration and frequency of administration) permit tumor elimination without a cytokine storm. Our conceptual model is simple enough to permit an understanding of a broad range of treatment outcomes (tumor elimination or escape, cytokine storm or no storm), yet detailed enough to also allow us to explore how the success and safety of a treatment regimen depends on drug and patient-specific properties. Given that a precise definition of a cytokine storm remains elusive, we also evaluate model predictions using different quantifiable definitions of cytokine storm. We end by outlining possible experimental approaches for parametrizing the model to increase its practical utility.

## Model Description

We propose the following system of five differential equations to capture the qualitative aspects of the dynamic relationship between tumor cells *T*(*t*), immune agonist *D*(*t*), immune cells *x*(*t*), and two types of cytokines *y*(*t*) and *z*(*t*) that regulate immune activity and act synergistically during a hyperactive immune response (22):

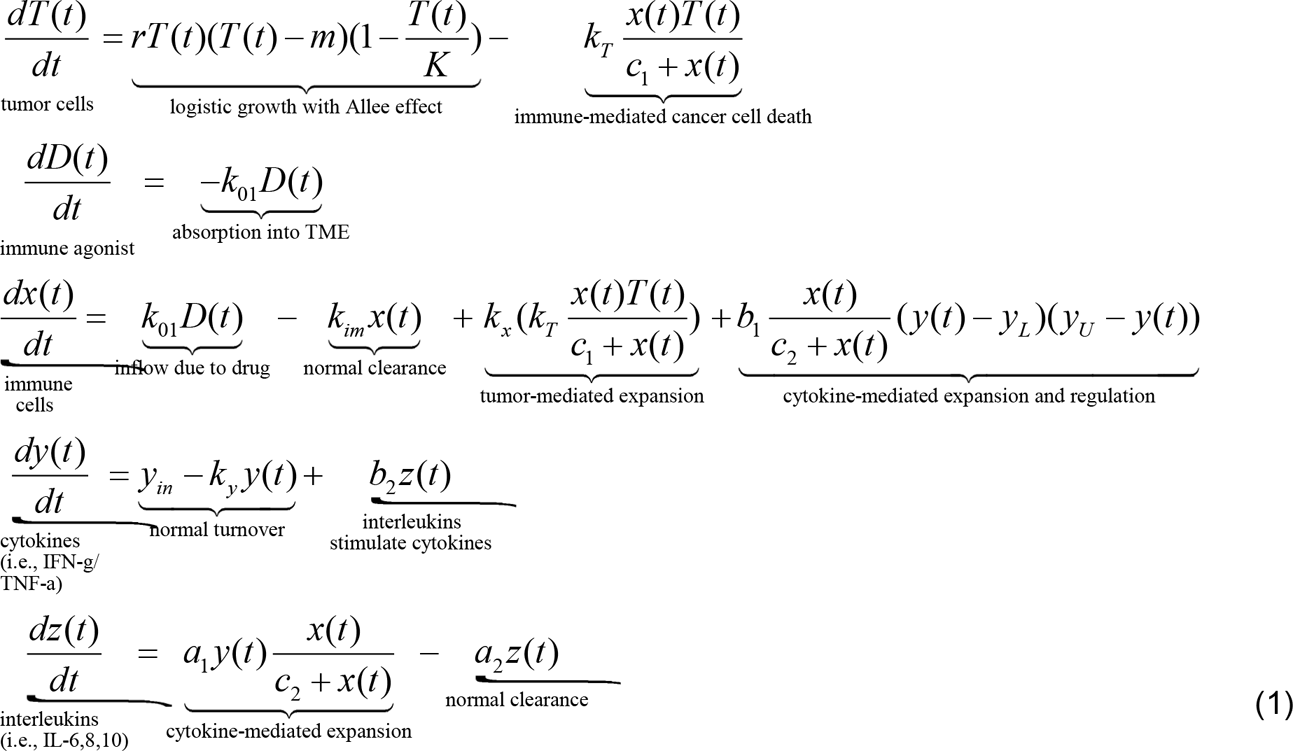

In this system, we assume that the tumor grows at a rate *r* and that there exists a strong Allee effect (23) with a threshold value *m*>0, such that when tumor size falls below *m*, it tends to zero, while if the population size is above threshold *m*, it grows to the carrying capacity *K*. The inability of sufficiently small tumors to grow can be explained by the tumor not yet accumulating sufficient mutations, or because of immune system control. As an example, the growth kinetics of BT-474 luminal B breast cancer cells was shown to be best-described by a model structure that considers the Allee effect (24).

Cancer cells can be eliminated by immune cells *x*(*t*) according to the Michaelis-Menten term 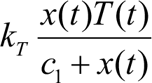, where *c*_1_ is the immune level for which the rate of tumor-killing is at half its maximum value, and *k_T_* is the maximal rate of cancer cell elimination by immune cells. The model assumes that the drug *D*(*t*) is administered exogenously and enters the tumor microenvironment at a rate *k*_01_*D*(*t*). As the drug is an immune agonist (like CAR T), it becomes available as a source term of anti-tumor immune cells *x*(*t*). Immune cells are further assumed to have a natural death rate *k_im_*. The immune cell population can additionally increase at a rate proportional to the rate of immune-induced tumor killing, as described by the term 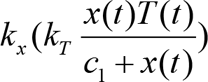. Finally, immune expansion is also regulated by the cytokines *y(t)* as follows: we assume that there exists a range of values for *y*(*t*) that act in an immune stimulatory fashion, but at a certain level beyond that they become immune inhibitory. In particular, the immune cells have an additional growth term when the concentration of cytokines is between *y_L_* _<_*_y_*_(*t*) <_*_yU_*, thereby capturing in a phenomenological way the dual stimulatory and inhibitory property of cytokines on the immune system (25,26).

Next, we describe the dynamics of cytokines *y*(*t*), which are involved in direct regulation of cytotoxic T cells *x*(*t*); these can be interpreted as TNF-alpha or IFN-gamma (27). We assume that these cytokines have a normal turnover rate and thus maintain an infection-free baseline level 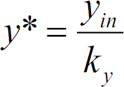. Further, at rate *b*_2_, these cytokines can be stimulated by other molecules *z(t),* such as IL-6, that are indirect regulators of immune cells (28). Finally, we assume that interleukins *z*(*t*) are produced in response to interactions between immune cells *x*(*t*) and cytokines *y*(*t*) at a maximal rate of *a*_1_ and are cleared at a natural rate *a*_2_. A schematic representation of this model structure is given in Figure 1A.

**Figure 1.**
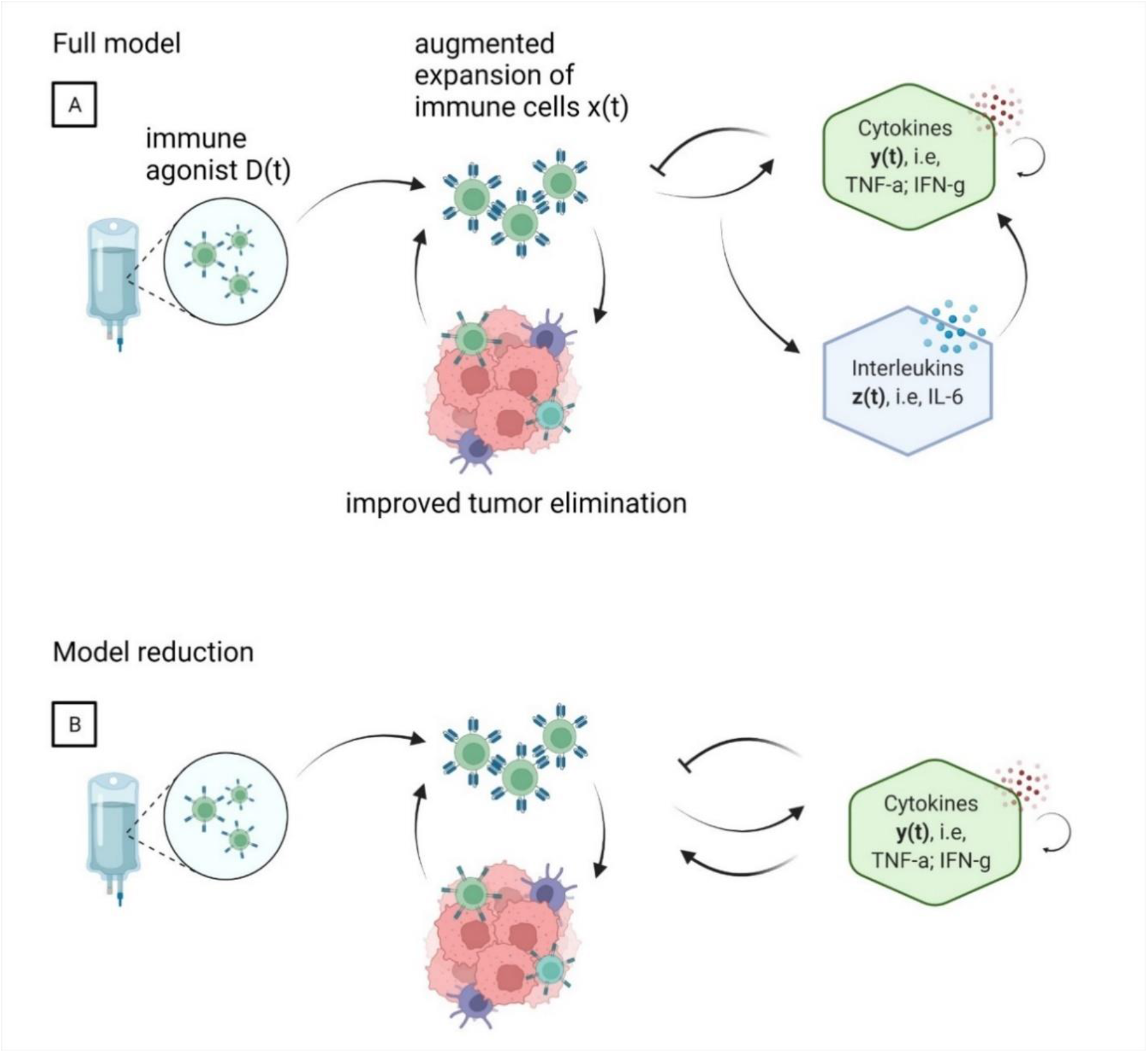
Schematic representation of immune-cytokine interactions subject to perturbation by an immune agonist *D*(*t*). (A) Full system as described by System (1) (B) Reduced schematic described by System (3).

Next, we assume that compared to the dynamics of the immune cells *x*(*t*), the *z*(*t*) subsystem reaches a quasi-steady state before it can affect immune cells *x*(*t*), since that half-lives of IFN-gamma and IL-6 are measured in hours (29,30), while tumor-immune dynamics typically take place on the scale of at least days. Taking 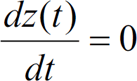 leads to interleukins *z*(*t*) reaching a quasi-steady state 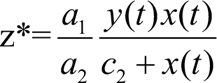. Substituting this expression into System (1), we get the following equation for the dynamics of cytokines:

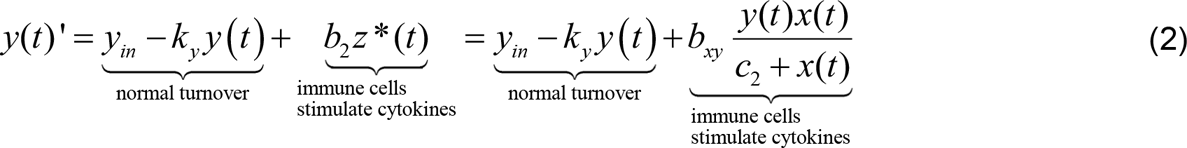

where 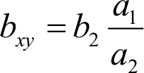 represents the rate at which immune cells stimulate cytokines. For notational consistency, with define *b_yx_* = *b*_1_ to represent the rate at which cytokines stimulate the immune cells. A schematic representation of this reduced system is shown in Figure 1B.

Under the quasi-steady state assumption, the final system of equations becomes

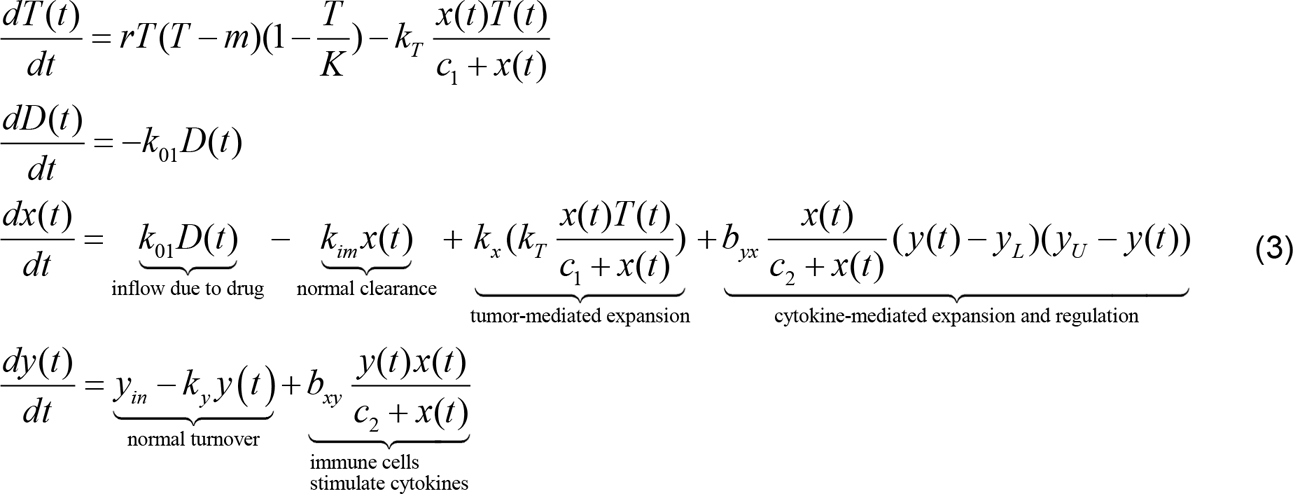

Given the phenomenological nature of the proposed model, parameter values were chosen arbitrarily to capture qualitatively different behaviors. Furthermore, since the model is not fit to specific data, units are chosen to be generic units of time and volume that can be specified when necessary for the purposes of a specific data set. A summary of default parameter values used in the simulations is given in Table 1.

**Table 1.** Parameters used in System (3). Parameter values were chosen arbitrarily to allow to capture qualitatively different behaviors.

| *Parameter* | *Description* | *Value* | *Units* |
| --- | --- | --- | --- |
| $D(0)$ | Initial volume of exogenous anti-tumor immune cells | varied | vol |
| $T(0)$ | Initial tumor volume | 0.1 | vol |
| $x(0)$ | Initial size of population of anti-tumor immune cells | 0.0 | vol |
| $y(0)$ | Initial size of population of cytokines regulating anti-tumor immune cells | 0.0 | vol |
| $k_{01}$ | Rate of inflow of exogenous immune cells | 0.1 | 1/time |
| $k_{im}$ | Rate of normal immune cell clearance | 0.099 | 1/time |
| $k_x$ | Rate of tumor-mediated immune cell expansion | 2 | 1/vol |
| $k_T$ | Rate of immune-mediated tumor kill | 0.05 | 1/time |
| $c_1$ | Michaelis constant ($x(t)$ level at which reaction rate is half of maximum value) | 1 | vol |
| $c_2$ | Michaelis constant | 1 | vol |
| $b_{yx}$ | Rate of cytokine mediated immune cell expansion | 0.01 | 1/(time*vol.<sup>3</sup>) |
| $b_{xy}$ | Rate of immune-mediated cytokine expansion | 0.8 | 1/time |
| $y_L$ | Cytokine-mediated threshold of immune cell expansion | 1 | vol |
| $y_U$ | Cytokine-mediated threshold of immune cell regulation | 6 | vol |
| $k_y$ | Normal cytokine decay rate | 0.42 | 1/time |
| $y_{in}$ | Cytokine production rate | $10^{-4}$ | vol/time |
| $r$ | Tumor proliferation rate | 0.06 | 1/time |
| $m$ | Allee threshold for tumor growth | 0.05 | vol |
| $K$ | Tumor carrying capacity | 1 | vol |

## Results

All simulations were conducted on MATLAB^®^ using ode23s as the numerical solver. The code is available at https://github.com/jgevertz/CytokineStorm.

### Cytokine storm can be decoupled from efficacy

We first evaluate whether the efficacy of exogenous immune agonists with respect to tumor elimination is necessarily coupled with a cytokine storm. Herein, we define a cytokine storm as cytokine levels *y*(*t*) surpassing the upper threshold *y_U_* at any point in time after treatment. In our simulations, we let the tumor grow until *t* = 200, and then simulate administration of either single or multiple doses of the drug described by *D*(*t*).

We start by confirming that lack of drug results in unrestrained tumor growth up to carrying capacity (Figure 2, column A) without any immune effects on its growth trajectory. Next, we simulate administration of a single dose of a drug, much as would be done with a CAR T cell therapy, at an arbitrary dose of 10 units (Figure 2, column B), 20 units (Figure 2, column C) and 25 units (Figure 2, column D). As can be seen in Figure 2B, the low dose of 10 units of the drug does not sufficiently stimulate the immune system, resulting in treatment failure without a cytokine storm. Figures 2C, D (row 1) numerically demonstrate that doses of 20 units and higher result in tumor elimination. This numerical result can be confirmed analytically by observing that the steady state (*T*^∗^, *D*^∗^, *x*^∗^, *y*^∗^) = (0,0,0, *y*_*in*_/*k*_*y*_) is stable and that any solution that passes through the point 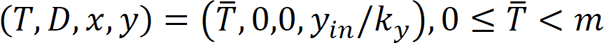 is attracted to this stable steady state. As doses of 20 and higher all (effectively) satisfy this property, we can be guaranteed that these solutions monotonically decay to zero and that relapse at a later time point is not possible for these doses. Next, we observe that tumor elimination may or may not be accompanied by a cytokine storm, as the dose of 20 units does not cause a storm (Figure 2C, row 4), but the dose of 25 units causes a storm (Figure 2D, row 4). This qualitative simulation suggests that a single dose of the drug can achieve efficacy without causing a cytokine storm, but the range of doses needs to be carefully selected to decouple efficacy from a storm.

**Figure 2.**
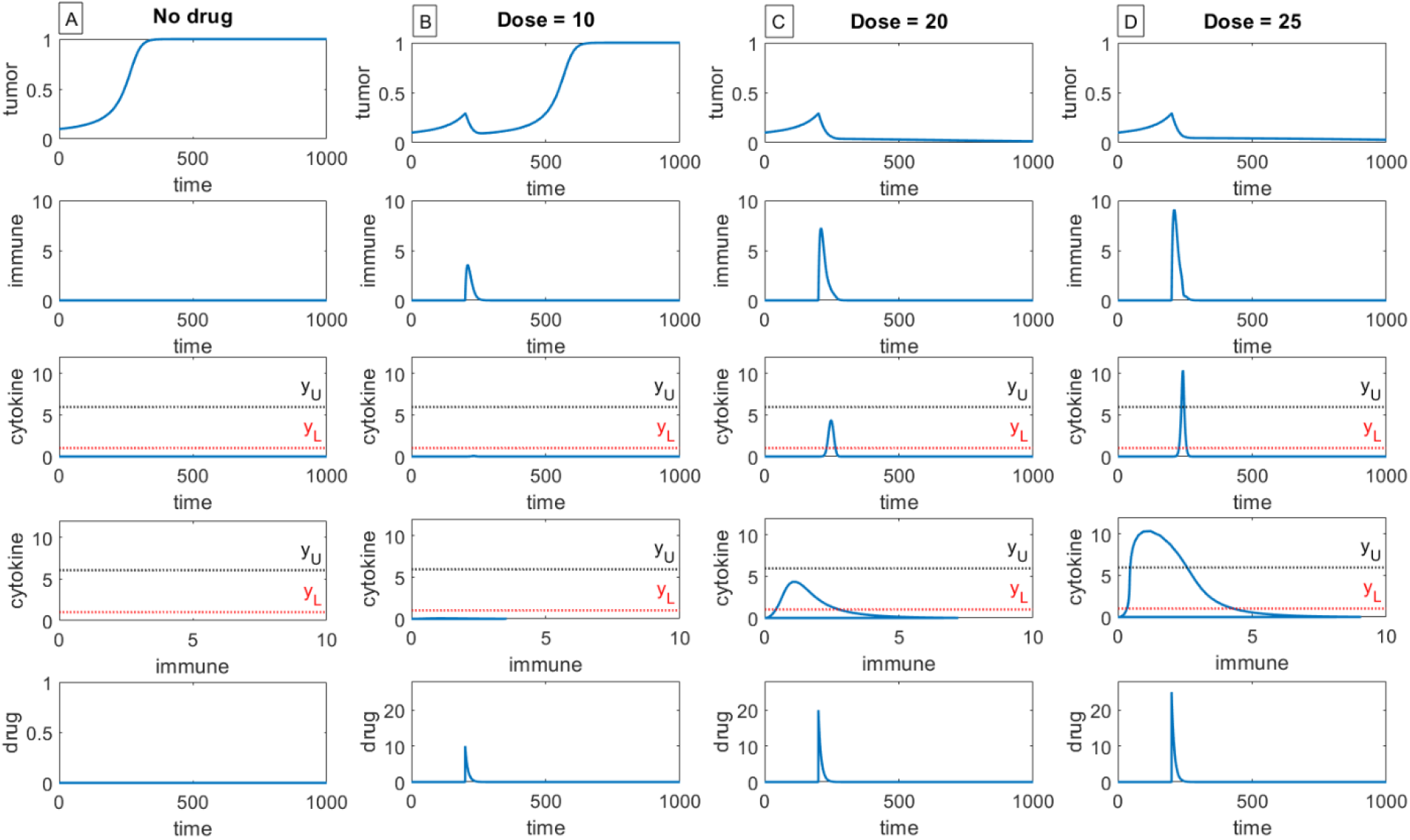
Model simulations in the case of (A) no drug, (B) a single dose of 10 units, (C) a single dose of 20 units, (D) a single dose of 25 units. First row shows tumor dynamics, second row shows immune dynamics, third row shows cytokine dynamics, fourth row shows dynamics in the immune-cytokine plane, and fifth row shows drug dynamics. A cytokine storm can be visually identified on the immune-cytokine plot in row 4 as a “loop” that surpasses threshold *y_U_* on the cytokine axis. We observe that single dose administration of the drug can achieve efficacy without causing a cytokine storm, but the magnitude of the dose must be carefully monitored in order to decouple efficacy from cytokine storm.

In these simulations, we define a cytokine storm as cytokine levels surpassing their upper threshold *y*_*U*._ Another way to visualize such a storm can be identified in the phase portraits in row 4 of Figure 2, where cytokines are plotted against immune cells. In this space, a cytokine storm looks like a hysteresis loop whose peak surpasses threshold *y_U_* in the cytokine axis. In a subsequent section, we will explore alternative definitions for a cytokine storm.

Notably, there exist regimes where insufficiently high dose can result in transient tumor suppression, followed up by a relapse (Figure 3). Within the frameworks of this model, relapse occurs due to insufficiently high dose. Together with Figure 2, these results highlight the importance of finding a sufficiently high dose to achieve efficacy but keeping it low enough to prevent a cytokine storm.

**Figure 3.**
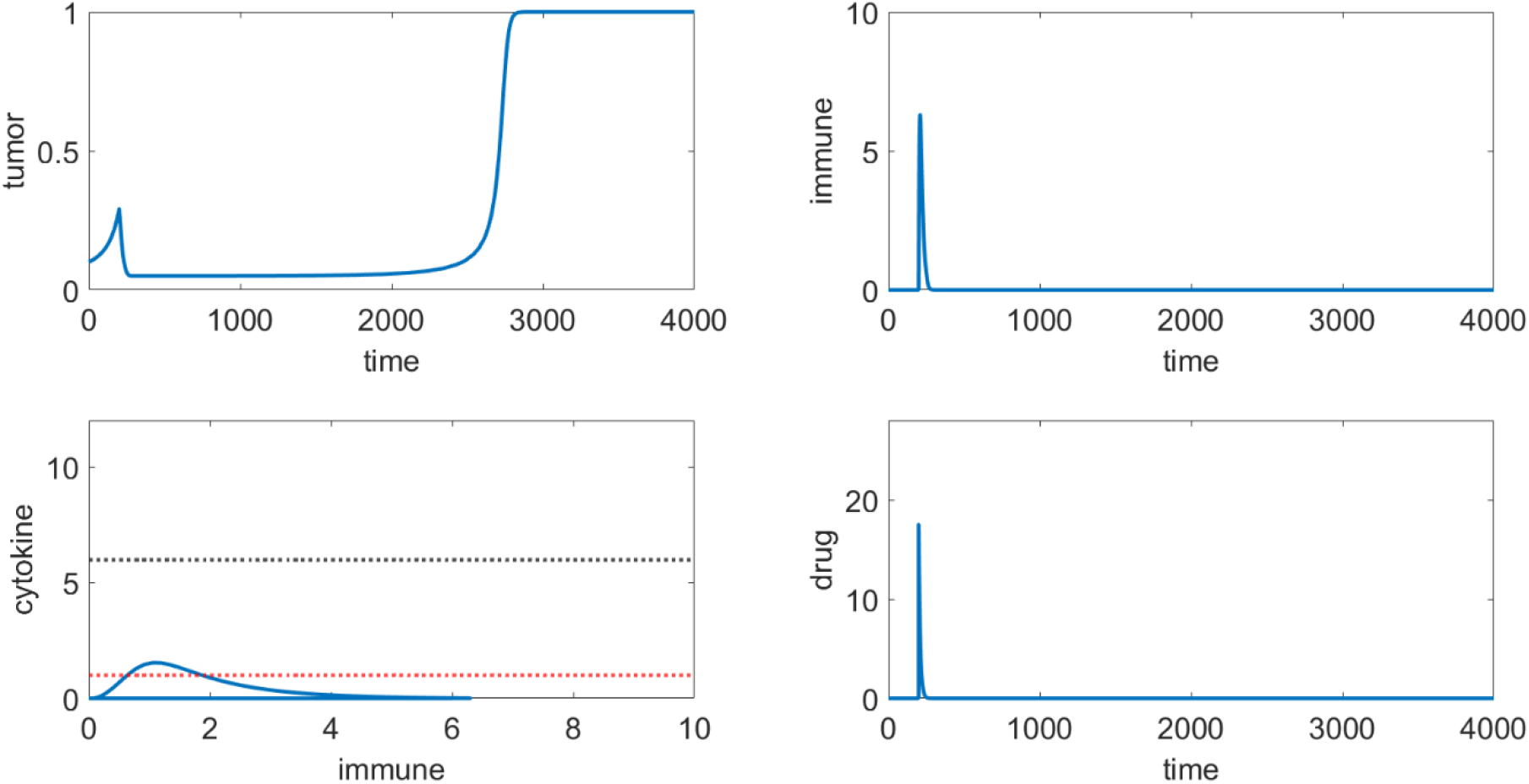
Model simulations of administering a single dose of 17.5 units. The model predicts transient tumor suppression by the immune system followed by a relapse at *t* = 2500.

### Low dose administration can achieve storm-free efficacy with sufficient number of doses

Next, we evaluate if the same level of efficacy can be achieved using lower drug doses in order to mitigate the likelihood of cytokine storm. For that, we simulate administration of 10 units of the drug, given as a single dose, or as two doses administered either 336, 168 or 24 time units apart (Figure 4). As can be seen in Figure 4C, adding a second dose can achieve efficacy without a cytokine storm if the two doses are given at an appropriately spaced time interval; spacing the doses out too much (Figure 4B) does not improve efficacy over a single dose, while administering them too closely together (Figure 4D) can result in a cytokine storm (row 3 showing cytokine levels, and row 4 showing a “storm loop”).

**Figure 4.**
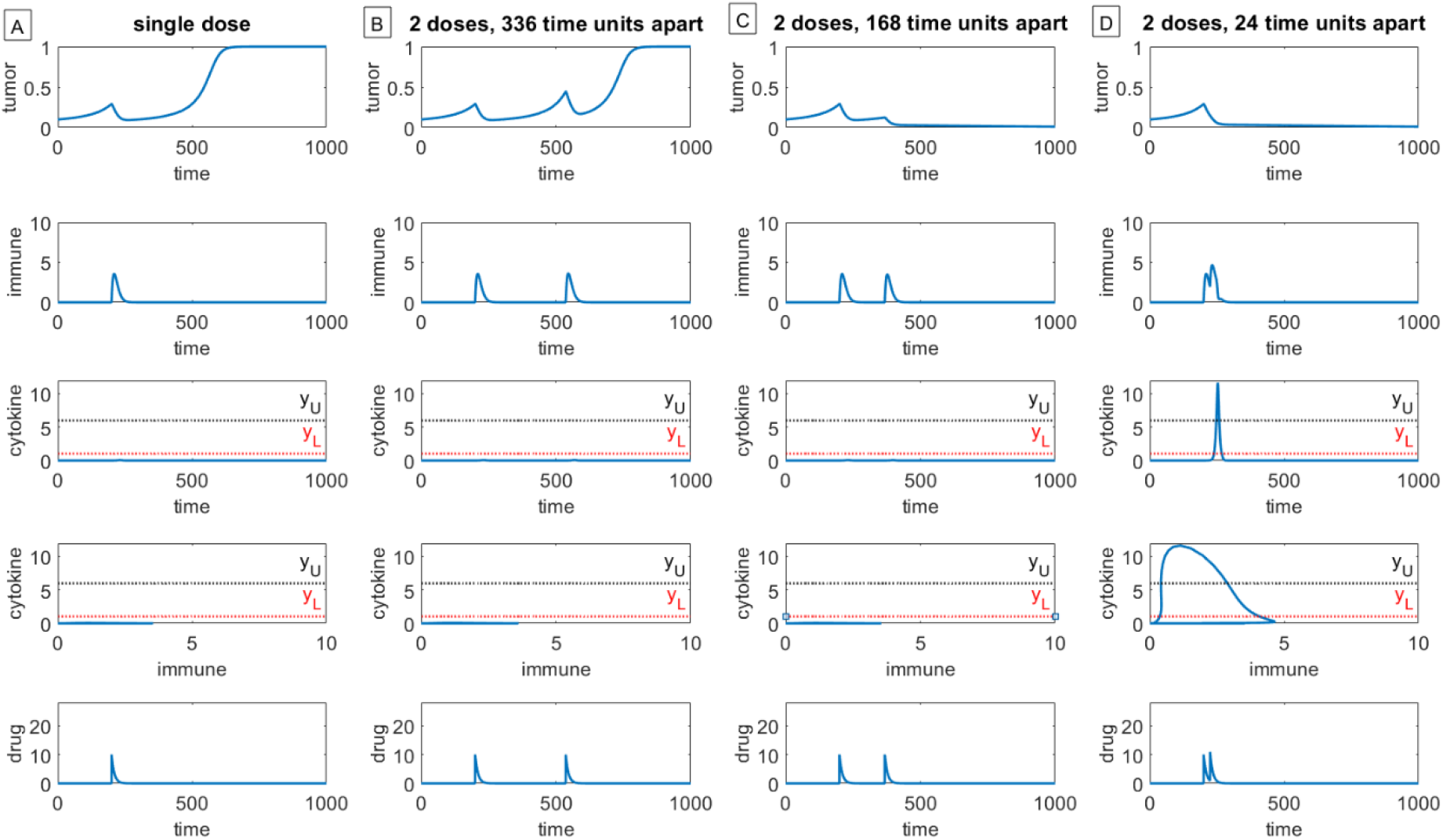
Impact of dose frequency on efficacy. Model simulations of 10 units of drug given (A) as a single dose, (B) as two doses given 336 time units apart, (C) as two doses given 168 time units apart, and (D) as 2 doses given 24 time units apart. First row shows tumor dynamics, second row shows immune dynamics, third row shows cytokine dynamics, fourth row shows dynamics in the immune-cytokine plane, and fifth row shows drug dynamics. Low doses can be efficacious without cytokine storm if administered at an appropriate frequency.

### Importance of initial tumor size for efficacy

In the previous set of simulations, we initiated treatment at *t* = 200, when the simulated tumor reached a size of 0.29 (29% of its carrying capacity). Here we simulate administration of a dosing protocol that proved efficacious for a tumor of this size and explore what happens if treatment is initiated a different time (meaning, when the tumor is at a different size at the onset of treatment). Specifically, in Figure 5A and Figure 5B we simulate administration of a 10 unit dose, given 2 times, where each dose is spaced out by 168 time units, with treatment administration starting either at *t* = 200 or *t* = 250, respectively. We observe that the same treatment is efficacious for a smaller tumor (columns A in Figure 5) but not for a larger one (column B in Figure 5).

**Figure 5.**
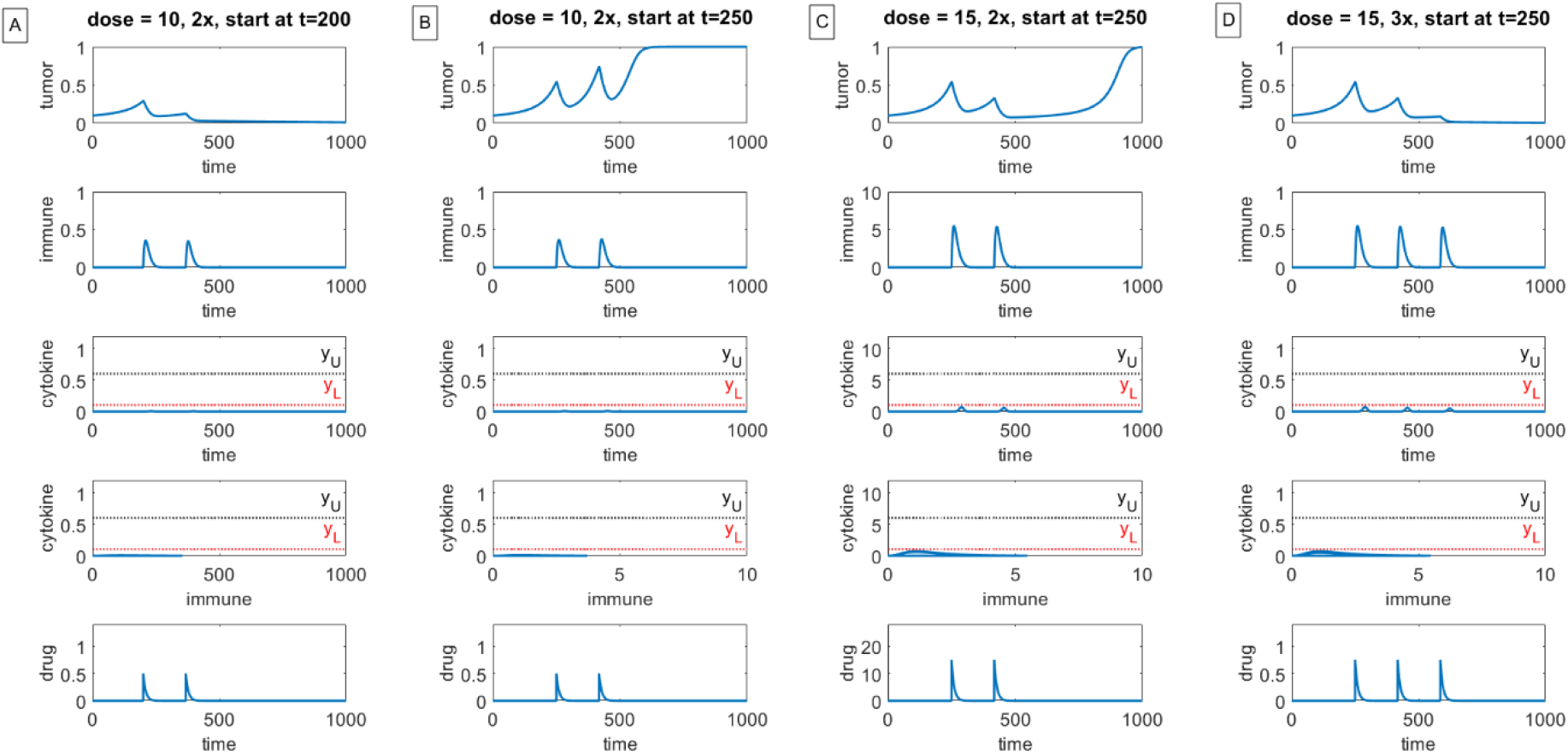
Impact of initial tumor size on treatment efficacy. Two doses of 10 units can (A) result in elimination if treatment is initiated at *t* = 200 but (B) not when treatment is initiated at *t* = 250 and is thus larger than if it were treated earlier. (C) Increasing the dose to 15 units at treatment initiation at *t* = 250 is insufficient to result in efficacy, but (D) adding a third dose of 15 units can achieve efficacy without a storm.

Next, we asked what could be done to control tumors that are larger at the time of treatment initiation. We first tried increasing the dose to 15 units administered every 336 time units but found that if treatment starts at *t* = 250, this increased dose is not sufficient to treat the tumor (Figure 5C). However, a third administration of the drug, given 168 time units later, can achieve tumor remission without a cytokine storm (Figure 5D). This emphasizes that the choice of protocol should depend on the size at the tumor at the initiation of treatment, and that there is unlikely to be a one-size-fits-all dosing protocol.

Given sensitivity of the projected treatment efficacy to initial conditions, a natural follow-up question is: how does the dose required to prevent additional tumor growth depend on the tumor size at the start of treatment? A critical advantage of a conceptual mathematical model is the ability to explicitly calculate a so-called “tumor static concentration”, or the “tumor stasis dose” (TSD) (31,32), which is a dose that is sufficient to prevent additional tumor growth but not to eliminate it. Here, for a fixed tumor size *T_s_* at the time of treatment initiation, tumor stasis requires:

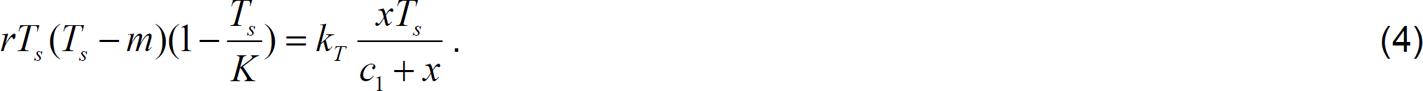

If we define 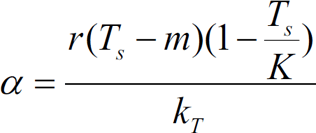, then

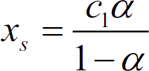

is the minimally necessary volume of immune cells needed to prevent the additional growth of a tumor with volume *T*_s_ at the time of treatment initiation. From here, we can further calculate the drug volume that is needed to achieve this *x*_s_. Given that treatment-induced cytokine storm in CAR T cell therapy typically occurs within a week of administration (33–35), it is reasonable to assume that the immune-cytokine dynamics occur on a fast time scale relative to tumor-immune interactions, and can thus be approximated to be at a quasi-steady state at the start of treatment. Under this assumption, if we define

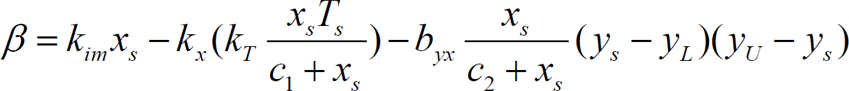

then the TSD can be calculated as

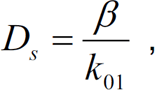

where 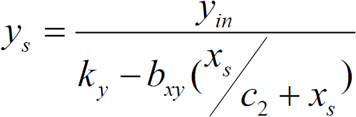 and *T_s_* and *x_s_* are defined above.

Such a basic calculation may already provide a framework to estimate the range of minimally necessary doses and schedules, given the tumor volume at the time of treatment initiation. That said, this calculation also requires knowledge of several basic patient-specific parameters, the impact of which is discussed in the next section.

### Sensitivity and variability analysis

In the previous section, we showed that a single high dose of the drug is likely to result in tumor elimination, which is important for therapies such as CAR T that are typically administered as a single dose. However, a high dose carries a serious risk of a cytokine storm. This has also been observed clinically, including in a single center phase 1-2a study of the CAR T cell medication tisagenlecleucel (ClinicalTrials.gov number NCT02435849). In this study, 75 patients received a single infusion of tisagenlecleucel, and high initial response rates were observed. In particular, rates of relapse-free survival and overall survival at six months were 73% and 90%, respectively, and 50% and 76% at twelve months. However, 77% of patients developed a cytokine storm, including both patients who eventually relapsed and those who did not (12). As another example, in (13) the authors evaluated 53 adults with acute lymphoblastic leukemia (ALL) who received 19-28z CAR T cell; while complete remission was observed in 83% if the patients, a cytokine storm occurred in 14 out of 53 patients (26%), with patients with higher disease burden exhibiting a greater incidence of cytokine storm and a lower likelihood of long term survival as compared to patients with a lower disease burden.

While our goal here is not to quantify the dose-response relationships reported in these cases, we can explore whether dose fractionation strategies may allow achieving efficacy without cytokine storm. It is also notable that currently manufacturing of CAR T cells can be logistically challenging (36,37), and therefore finding ways to minimize the number of necessary doses to achieve efficacy without cytokine storm is warranted.

For that, we evaluate what intrinsic properties of both the drug and the patient, as defined within the frameworks of the proposed model, can affect efficacy and safety. We start by performing a one at a time (OAT) local sensitivity analysis on all model parameters, only excluding the Allee threshold and carrying capacity in the tumor growth term as these are unlikely to be accurately estimated for each individual patient. This analysis quantifies how a ±*N*% change in a single baseline parameter value (as defined in Table 1) changes the output in both the tumor volume at *t* = 400 (200 time units post treatment initiation), and the maximum cytokine value over the 400 unit window of time. We considered 1 ≤ *N* ≤ 50, meaning parameters were perturbed as little as 1%, and as much at 50%, from their baseline value. The dosing strategy used was to administer a single dose of 20 units of drug, starting at *t* = 200. This was chosen because, as we saw in Figure 2, this dose results in tumor eradication without a storm. Administering a lower dose is not an effective treatment and administering a higher dose results in a cytokine storm.

The outcomes of this OAT local sensitivity analysis for both a 1% and a 50% variation in the baseline value of each parameter are found in Figure 6. We learn that the rate of cytokine stimulation by the immune system *b_xy_* and the natural turnover rate of cytokines *k_y_* are two of the most sensitive model parameters on both tumor volume and the maximum cytokine level. *b_xy_* is the most sensitive parameter on both outputs when parameters are varied by 1%, and it is also the most sensitive parameter affecting the maximum cytokine level when it is varied by 50%. It is the fourth most sensitive parameter on tumor volume when varied by 50%. Whether it is varied by 1% or 50%, *k_y_* is the second most sensitive parameter affecting the maximum cytokine level, and the third most sensitive parameter affecting the tumor volume. Not surprisingly, the tumor growth rate *r* and the drug-induced tumor death rate *k_T_* are both sensitive parameters for tumor volume (ranking in the top four independent of the percent variation considered), though the sensitivity of each of these parameters is significantly diminished when considering their impact on the maximum cytokine level and thus on the likelihood of developing a cytokine storm.

**Figure 6.**
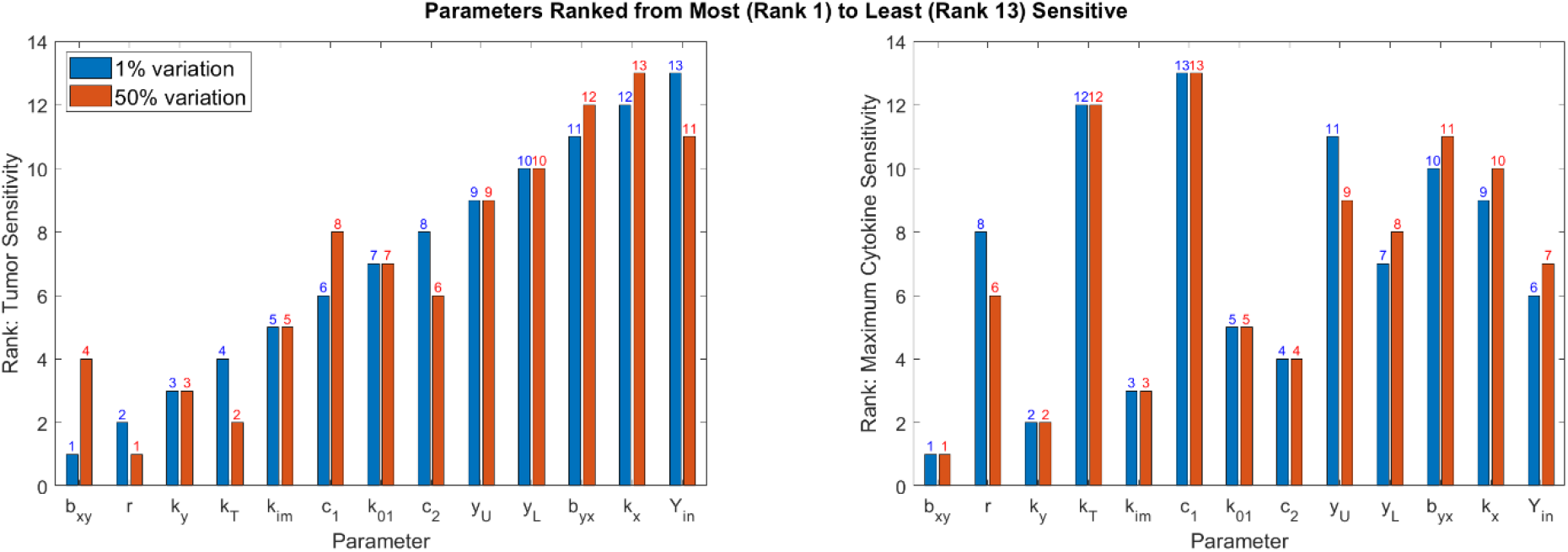
Local sensitivity analysis. Model parameters are perturbed, one at a time either up or down 1% (blue) to 50% (red) from the baseline value presented in Table 1. Parameters are then ranked by the impact they have on a specified output: (left) impact on tumor volume at *t* = 400, and (right) impact on maximum cytokine level over 0 ≤ *t* ≤ 400. A rank of 1 indicates the parameter is the most sensitive for a given perturbation percent and output, whereas a rank of 13 indicates the parameter is the least sensitive to a given perturbation and output.

### All four regimes can occur for the same dose depending on individual’s characteristics

This analysis led us to further explore the impact of the immune stimulation rate of cytokines (*b_xy_*) and the rate of tumor killing (*k_T_*) on both treatment efficacy and toxicity (i.e., the development of a cytokine storm). These sensitive parameters represent key features of the drug and the tumor, as compared to a sensitive parameter like *k_y_*, which represents a natural turnover rate of immune cells, but is not dependent on the drug or the treatment. To explore the impact of these parameters on treatment outcome, we allow *k_T_* to vary over the range [0.004, 0.164] and we vary *b_xy_* over the range [0.6, 1.55]. All other parameters are fixed as defined in Table 1. The model is solved over a set of 121×121 parameter values that are uniformly spaced over the specified range, using a single dose of 10 units administered at time *t* = 200. For each parameter set, we classify the treatment as being in one of four regimes, as summarized in Table 2.

**Table 2.** Four potential outcomes for tumor response to treatment with immune agonist.

|  |  |
| --- | --- |
| Efficacy + no storm<br>(regime P1) | Efficacy + storm<br>(regime P2) |
| No efficacy + no storm<br>(regime P3) | No efficacy + storm<br>(regime P4) |

In the results that follow, we define a cytokine storm as having occurred if the cytokine levels surpass the upper threshold of *y_U_*, and we define a treatment to be effective if the final tumor volume is at least 90% below its carrying capacity (i.e., *T*(3600) ≤ 0.1*K*). Figure 7 demonstrates that all potential outcomes described in Table 2 are achievable for this fixed protocol (10 units of drug administered at t = 200), depending on the value of the immune stimulation rate of cytokines and the rate of tumor killing. At low values of cytokine stimulation and tumor killing, the model predicts that the outcome of treatment falls in regime P3, with no tumor killing and no cytokine storm. Sufficiently increasing the rate of killing can push the model into regime P1, where the tumor is effectively treated without a cytokine storm. On the other hand, sufficiently increasing the rate of cytokine stimulation can result in a cytokine storm without tumor elimination (the worst-case scenario, regime P4). Sufficiently increasing both parameters pushes the model into regime P2, in which case an effective treatment is accompanied by a cytokine storm. Though not particularly surprising, we also observe that for this treatment protocol, whether a storm occurs is much more sensitive to the rate of cytokine stimulation *b_xy_* than to the rate of tumor killing *k_T_*.

**Figure 7.**
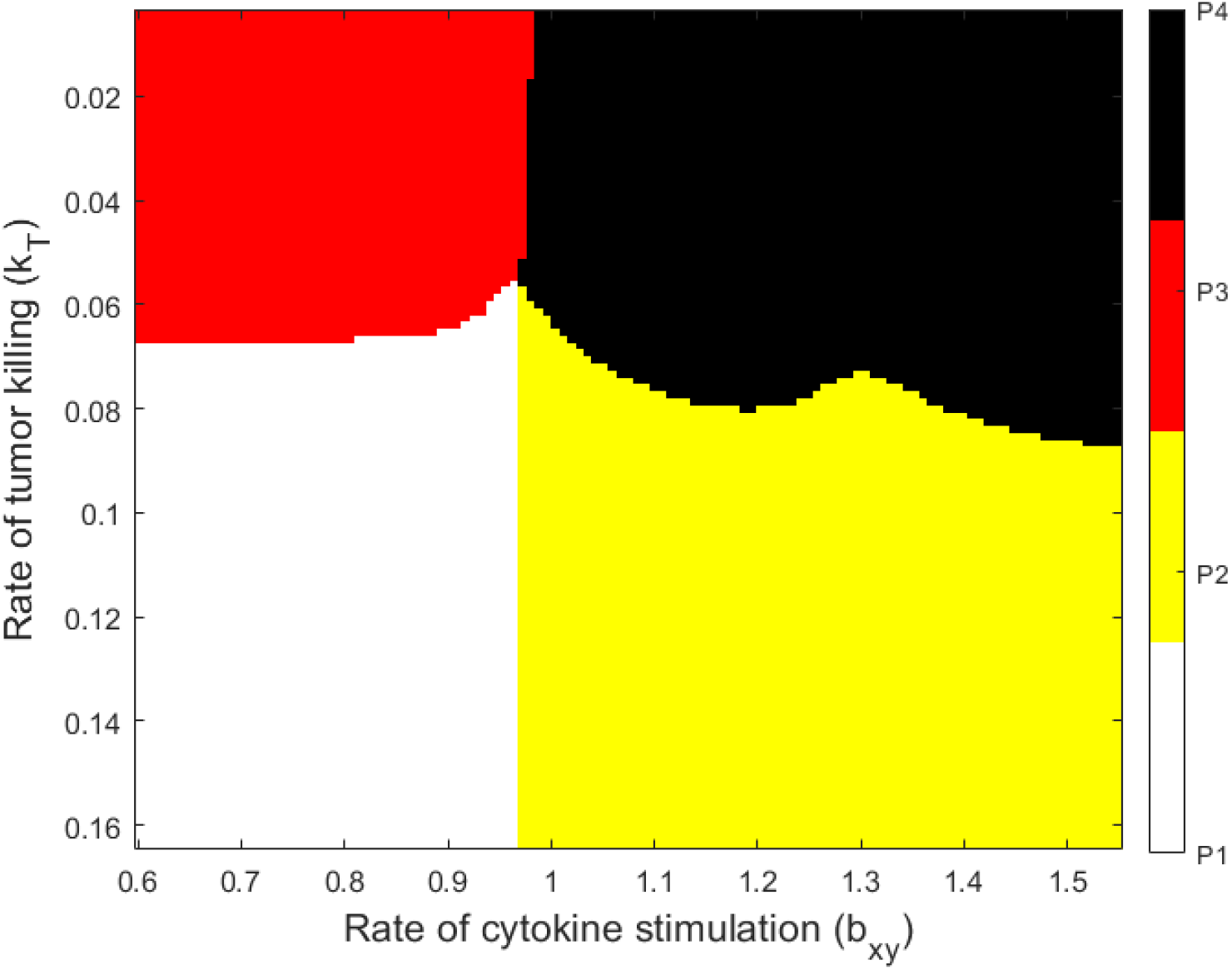
Treatment efficacy and toxicity as a function of rate of cytokine stimulation (*b_xy_*) and rate of tumor killing (*k_T_*) in response to a single administration of the drug at dose 10 at *t* = 200. Each parameter set is classified as falling into one of four parameter regimes. P1 (white): effective treatment with no cytokine storm (best case). P2 (yellow): effective treatment but cytokine storm occurs. P3 (red): ineffective treatment, no cytokine storm. P4 (black): ineffective treatment with a cytokine storm (worst case).

If we think of each point in this parameter space as representing a plausible “virtual patient” (38,39), we can immediately see a problem with treating everyone with a single dose of 10 units. While some individuals will be effectively treated with limited toxicity (no cytokine storm), other individuals run the risk of a cytokine storm accompanying the effective treatment, and others may not be able to be effectively treated by this dose, toxicity considerations aside. This phenomenon, where a therapy effective for one virtual patient has the potential of being dangerous for others, as has been seen in other studies involving virtual patients (40,41).

### Cytokine storm mitigation through dose modulation

Next, we consider a virtual patient for which a single dose of 10 units administered at *t* = 200 results in an effective treatment but causes a cytokine storm. We choose *b_xy_* = 1.55 and *k_T_* = 0.15, which falls in regime P2 in Figure 7. We ask whether it is possible to control the storm but preserve efficacy at the fixed cumulative dose of 10 units through dose fractionation (administering the same total dose but divided over different time intervals to mitigate toxicity). To test this, we take the effective dose of 10, divided up into two to sixteen doses, spaced out over 24-192 time units. For instance, the first column in Figure 8 shows the output of giving two doses of drug, meaning each dose is 5 units. Across each row, we look at what happens if those two doses are spaced out anywhere from 24-192 time units apart.

**Figure 8.**
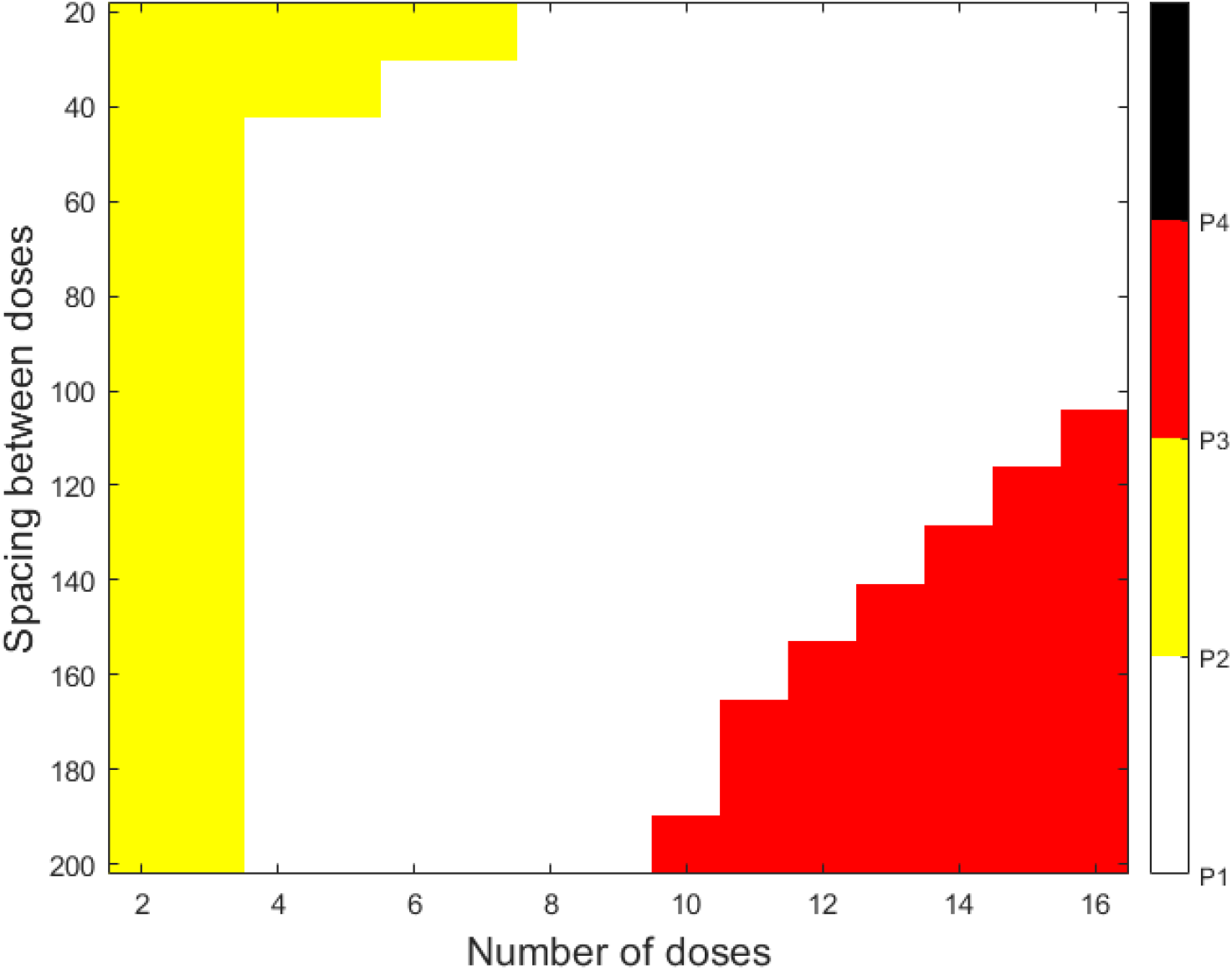
Treatment efficacy and toxicity as a function of the number of doses and the spacing between doses, when the cumulative dose given is 10 units. Results are for *b_xy_* = 1.55 and *k_T_* = 0.15, a parameter set for which the default treatment of one dose of 10 is effective but causes a cytokine storm. Each treatment strategy is classified as falling into one of four parameter regimes. P1 (white): effective treatment with no cytokine storm (best case). P2 (yellow): effective treatment but cytokine storm occurs. P3 (red): ineffective treatment, no cytokine storm. P4 (black): ineffective treatment with a cytokine storm (worst case).

As can be seen in Figure 8, there exists a “sweet spot” in the dosing space, where the properly selected dose fractionation strategy can preserve efficacy without a cytokine storm. If the number of doses is too small, and therefore the amount of drug given at each dose is too high, no fractionation strategy can prevent the cytokine storm (see two and three doses in Figure 8). At an intermediate number of doses, and therefore an intermediate amount of drug administered at each treatment, there exist a range of fractionation schedules that can effectively eradicate the tumor without a cytokine storm. As the number of doses increases (i.e., the amount administered per dose is quite low), the concern shifts from a possibility of a cytokine storm to limited efficacy of the treatment. If the doses are sufficiently close together, the treatment can be effective without a storm. But, if the doses are spread out too much, treatment is likely not to be effective (see doses twelve, fourteen and sixteen in Figure 8).

Notably, three of the four regimes can be observed through dose fractionation, further highlighting that a drug can be both efficacious and detrimental, and that understanding of several key characteristics specific to an individual patient could determine the “sweet spot” and therefore the outcome of their treatment. Furthermore, while there exist calculated boundaries for each “treatment sweet spot”, the protocol is most likely to be both safe and efficacious if chosen in the middle of such a “sweet spot”. As such, even if assessment of patient-specific parameters are not perfectly accurate, the projected output is robust to small perturbations and is therefore likely to still achieve both safety and efficacy.

### Model insensitivity to cytokine storm definition

Despite its importance, a precise quantifiable definition of cytokine storm remains elusive. For instance, Giavridis et al. (42) focus on absolute levels of cytokines as a strong correlate of cytokine storm severity and survival in murine models. On the other hand, Davila et al. (43) focus on fold change of cytokines rather than absolute values as markers of cytokine storm severity, and Lee et al. (28) suggest that a relative rate of change of cytokine levels may be a more appropriate marker. To discriminate between the differences in these definitions of a treatment-induced cytokine storm, we conducted simulations by defining cytokine storm as either based on absolute levels of cytokines (as used throughout the manuscript), or on the rate of change of cytokine expansion. Interestingly, within the frameworks of the proposed model, the projected cytokine storm occurrence is essentially invariant to either of these definitions, provided the cutoff for a storm in the rate of change case is reasonably defined (we use a cutoff rate of 0.417). In particular, only one of 121×121 = 14,641 parametrizations considered in Figure 7 changes classification under the rate of change definition of a storm: 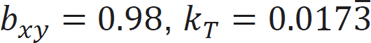 is categorized as P4 (no efficacy + storm) using absolute cytokine levels, but P3 (no efficacy + no storm) using the rate of change in cytokine levels.

## Discussion

Here we proposed a conceptual mathematical model of immune-cytokine interactions capable of describing how treatment with an immune cell agonist may result in the transition from a normal immune response to a response that can be interpreted as a cytokine storm. The goal of the model was not to describe a particular data set or to incorporate extensive biological detail, but to capture qualitative relationships between the broad classes of immune cells and cytokines that are sufficient to reproduce these dynamics. We further sought to identify key parameters that may suggest whether an individual may be susceptible to experiencing a cytokine storm.

For this purpose, we proposed a five-dimensional ordinary differential equation model, reduced to four dimensions using a quasi-steady state argument, that describes the dynamics of cancer cells, immune cells, cytokines, and a drug that directly increases the population of immune cells. The model revealed the existence of four possible regimes (Table 2, Figure 7, 9): treatment efficacy without a storm, efficacy with a storm, no efficacy without a storm, and efficacy with a storm. We sought to explore the plausibility of being in the first parameter regime (effective treatment with no storm) as a function of model parameters and dosing schedules.

**Figure 9.**
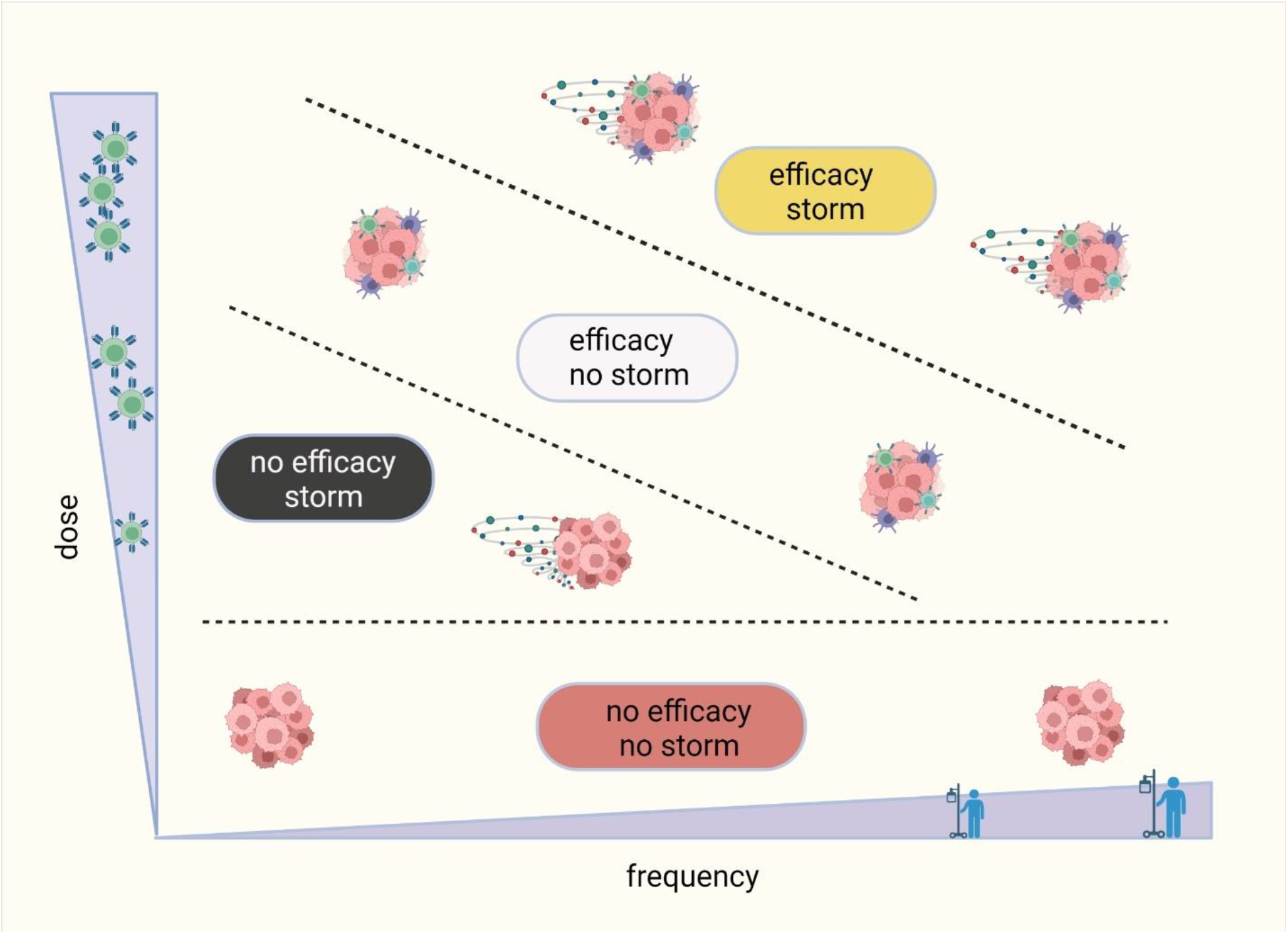
Dose-frequency tradeoff for maintaining anti-tumor efficacy while mitigating the risk of cytokine storm.

Simulations revealed that a tumor can be eliminated without a cytokine storm using both high and low dose regimens. As expected, high dose regimens carry a greater risk of generating a cytokine storm, and careful dose planning and monitoring is required to prevent it. For low dose therapy, the larger concern is sufficiently activating the immune system to trigger tumor eradication. However, once the dose has been given often enough to be effective, the risk of a cytokine storm is much lower than at high doses. We note that these are qualitative conclusions of the model – they are not meant to make particular dosing recommendations but are instead guidelines of what to consider when treating with an immune cell agonist over a range of possible doses.

Our model further allowed us to explore if the emergence of a storm could be prevented with a change in dosing strategy. To this end, we considered a model parametrization classified as “effective treatment with a cytokine storm” when treated with a single high dose of drug. We then asked if it is theoretically possible to dose the drug differently (giving the same cumulative dose) to preserve efficacy while eliminating the cytokine storm. By varying the number of doses and spacing between them, we uncovered that tumor control without a storm can be achieved at an intermediate number of doses provided the doses are not spaced too far apart (which could result in loss of tumor control) but are also not too close together (which still results in a storm). While again this is not meant as a quantitative recommendation given that the model is not parametrized to treatment response data, this does highlight that it is theoretically possible to decouple treatment efficacy and storm occurrence.

Finally, we evaluated the sensitivity of model predictions to two possible quantifiable definitions of a cytokine storm, namely, one defined based on absolute number of cytokines (42) or one based on their rate of expansion (28). Interestingly, we observed that within the frameworks of the proposed model, the results are effectively invariant to the definition of the storm. This suggests that the lack of clarity surrounding the precise definition of a cytokine storm need not be an issue in studying these storms and considering how to mitigate them.

### Patient-specific parameter estimations

The model presented herein was designed to study qualitative aspects of treatment with immune cell agonists. Our major focus was to determine if it is theoretically possible to decouple treatment efficacy and the occurrence of a cytokine storm, even when given at the same cumulative dose. While the model can answer this theoretical question, its potential utility is of course limited by the fact that it is not calibrated to patient-specific data. Here we discuss potential avenues for collecting the necessary data to approximate the value of the more sensitive model parameters.

The parameter *k_T_* (rate of immune-mediated tumor kill) could be estimated using in vitro cytotoxicity assays (44), a very commonly used experimental system in pre-clinical development of oncology drugs. The typical output of the in vitro cytotoxicity assay depicts concentration of the drug that causes an *N*% reduction in the cell viability of the treatment group as compared to the control after a fixed period of time from treatment initiation. A sample protocol to collect these data would involve seeding cancer cells in a 96-well cell culture plate for 24 hours prior to treatment. Cells would then be treated with, for instance, ten different serially diluted concentrations of a drug, in duplicate. Replicate plates of treated cells would be incubated for different periods of time (i.e., 1, 2, 5, 8 days, as appropriate). At the specified harvest day, cells would be treated with a reagent that allows measuring luminescence of viable cells, after which relative cell viability would be calculated as a percentage of untreated cells. For therapies involving immune agonists, such metrics are assessed using co-incubation of tumor and effector cells for different effector:tumor (E:T) ratios, such as was done in (45,46). Normal cytokine decay rate can typically be estimated according to well-established formulas for exponential decay, where k_y_=log(2)/(half-life). Similar calculation can be applied to any specific cytokine for which half-life is known.

Estimating patient-specific values of parameter *b_yx_* (rate of cytokine mediated immune cell expansion) might be achievable through similar co-incubation methods, where immune cells are exposed to various concentrations of different cytokines, and the longitudinal data on immune cell expansion or regulation is collected and quantified. Such an experiment may additionally allow quantifying patient-specific values of *y_L_* and *y_U_*. Similarly, to establish the patient-specific value of *b_xy_* (rate of immune-mediated cytokine expansion) it might be possible to expose a fixed concentration of cytokines to different concentrations of immune cells, recording and quantifying corresponding rates of expansion.

The patient-specific tumor growth rate *r* can also be assessed through either in vivo data on the change in tumor size prior to treatment, or through in vitro growth of the untreated cells. Conducting the proposed set of experiments, along with measuring the tumor size *T*(0) at the initiation of treatment, would provide more realistic estimates for the key parameters identified in this work. This would allow the qualitative conclusions of the analyses presented herein to be made quantitative. If such experiments could be conducted on an individual basis prior to treatment initiation, clinicians could be empowered to design a patient-specific protocol that maximizes efficacy while mitigating the risk of cytokine storm.

## Conclusions

We demonstrate how a low-dimensional phenomenological model of cytokine-immune interactions during treatment with an immune agonist can lead to a better conceptual understanding of treatment efficacy and toxicity. Though the model is unlikely to have direct practical utility in its current form, it does lay the groundwork for understanding the complex, nonlinear relationship between the tumor-killing and cytokine-stimulating features of an immune agonist like CAR T therapy. Just as economists, who, according to Nobel Prize winner Ester Duflo, when helping governments design new policies and regulations, have to adopt the practical situation-specific mindset of a plumber (51), so should modelers when increasingly taking part in designing treatment protocols. “Plumbers try to predict as well as possible what may work in the real world, mindful that tinkering and adjusting will be necessary since our models give us very little theoretical guidance on what (and how) details will matter… Economists should seriously engage with plumbing, in the interest of both society and our discipline.” Adopting a similar mindset of “modelers as plumbers” will invariably increase practical utility of mathematical modeling of disease and treatment.

## Author Contributions

IK and JG contributed equally to all stages of manuscript preparation.

